# Degradation of resistant α-1,4-glucan by vaginal *Gardnerella* species is associated with bacterial vaginosis

**DOI:** 10.1101/2023.01.28.526025

**Authors:** Rosanne Hertzberger, Sara Morselli, Sara Botschuijver, Lisa Himschoot, Leon Steenbergen, Sylvia Bruisten, Warren Lewis, Piet Cools, Remco Kort

## Abstract

This study investigates the degradation of resistant α-1,4-glucan by vaginal bacterial species, with a focus on *Gardnerella* spp., to elucidate its role in bacterial vaginosis (BV). The ability of *Gardnerella vaginalis, Gardnerella swidsinskii, Gardnerella leopoldii, Gardnerella piotii, Lactobacillus iners*, and *Lactobacillus crispatus* was assessed to metabolize an ungelatinized, labeled form of raw amylose, a degradation-resistant α-1,4-glucan. The enzymatic activity of these species was evaluated *in vitro*, and its association with BV was examined in vaginal swabs. *Gardnerella vaginalis, G. swidsinskii*, and *G. leopoldii* demonstrated the best ability to degrade resistant α-1,4-glucan *in vitro*. Unlike the cell-bound, S-layer-associated glycogen-degrading activity in *L. crispatus*, this α-glucosidase activity in *Gardnerella* was also extracellular, but not cell-bound and not repressed by glucose. Vaginal swabs showing high rates of resistant α-1,4-glucan degradation activity were associated with BV, particularly in the concurrent presence of *G. leopoldii, G. swidsinskii*, and *G. vaginalis*. These findings suggest a role of α-1,4-glucan degradation in BV pathogenesis mediated by *Gardnerella* species. The results indicate the potential of targeting bacterial amylase activity as therapeutic strategy for BV prevention and treatment.

## Introduction

The vaginal mucosa of reproductive aged women is rich in glycogen (Cruickshank, 1934; Mirmonsef et al., 2016). Through shedding and lysis of the superficial vaginal epithelial cells, glycogen is released in the vaginal lumen where it can serve as a carbohydrate source for bacteria colonizing the vagina, including *Lactobacillus crispatus* (Hertzberger et al., 2022; van der Veer et al., 2019). Glycogen levels were found to be associated with the vaginal microbiota composition (Mirmonsef et al., 2014). The more *Lactobacillus*-dominated vaginal microbiota is associated with higher levels of glycogen compared to the more diverse *Lactobacillus*-deprived microbiota with higher levels of fastidious anaerobes such as *Gardnerella, Fannyhessea* and *Prevotella* (Dols et al., 2016). The latter microbial dysbiotic state is also referred to as bacterial vaginosis (BV), and is a general risk factor for adverse reproductive and sexual health outcomes, such as HIV infection (Gosmann et al., 2017), HPV infection (Lin et al., 2020), subfertility (Vergaro et al., 2019), endometritis (Haggerty et al., 2004), pregnancy loss (R. G. Brown et al., 2018) and preterm birth (Gudnadottir et al., 2022).

*Gardnerella* is a hallmark genus in BV (10). The vaginal microbiota in women with BV has a much higher bacterial diversity compared to the healthy, *Lactobacillus-*dominated microbiota. Species within the *Gardnerella* genus can cause many of the clinical manifestations of BV, including exfoliation of the epithelial wall lining the vagina and thinning of the mucus leading to abnormal secretions (Gelber et al., 2008; Gilbert et al., 2013; Lewis et al., 2013). *Gardnerella* spp. differ in their ability to produce sialidase found in *G. piotii, G. pickettii* and in a subset of *G. vaginalis* strains (Robinson et al., 2019; Sousa et al., 2023).

Alpha-glucosidases are enzymes that catalyze the hydrolysis of α1-4 or α1-6 linkages between glucose units of α-glucans. In an environment with glycogen as the prime carbohydrate source, α-glucosidases of vaginal bacteria could play a central role in carbon and energy metabolism by extracellular degradation of host-associated glycogen, thereby releasing smaller maltodextrins for utilization in the surrounding environment. Previously, we have suggested a genetic locus for one of these α-glucosidases: the *L. crispatus* amylopullulanase (Dols et al., 2016; Hertzberger et al., 2022; van der Veer et al., 2019). Strikingly, we found a large number of *L. crispatus* laboratory strains (23%) to be deficient for this amylopullulanase and identified frequently occurring mutations in the genes of these strains as well as in databases of vaginal metagenomes (Hertzberger et al., 2022; Sycuro et al., 2023). This suggests that carbohydrate availability for some strains may depend on glycogen cleavage by other bacteria or host amylase as well.

Most studies characterizing the α-glucosidases of vaginal bacteria have used soluble starch or glycogen (Bhandari et al., 2023; Nunn et al., 2020; Spear et al., 2014). However, for the gut microbiome it was shown that more resistant structures of α-glucan allow to discriminate between carbohydrate-degrading enzymes of different bacterial taxa: Animals fed with degradation-resistant α-glucan (resulting from a denser structure and a lower degree of branching), show an increase of amylolytic species such as *Bifidobacterium* spp. (I. Brown et al., 1997; Sybille et al., 2013). These bacteria produce a type II amylase-pullulanase fusion protein capable of hydrolyzing resistant starch (Jung et al., 2020; Maier et al., 2017). A homologue of this enzyme was identified in *Gardnerella vaginalis*, expressed in *Escherichia coli*, and found to degrade amylose, pullulan, glycogen and starch (Jenkins et al., 2023). More recent research showed that 14 out of 15 tested *Gardnerella* strains harbor a full copy of this gene (Bhandari et al., 2023) and that isolates of four different *Gardnerella* species can transport and metabolize the small maltodextrins derived from glycogen breakdown (Bhandari & Hill, 2023).

Here we aimed to use resistant α-1,4-glucan to distinguish between bacterial glycosidases, thereby increasing our understanding of their contribution to glycogen breakdown in various microbial contexts. We show that *Gardnerella* species can be distinguished from vaginal lactobacilli by their ability to break down resistant α-1,4-glucan. The degradation of resistant α-1,4-glucan appears to be associated with vaginal dysbiosis.

## Methods

### Bacterial cultivation

For this study, we selected various strains isolated from vaginal samples. The isolation and characterization of the *Lactobacillus crispatus* amylopullulanase-sufficient strain RL10 and amylopullulanase-deficient strain RL09 were previously described (Dols et al., 2016; Hertzberger et al., 2022; van der Veer et al., 2019). Various *Gardnerella* strains were included as well as other members of the vaginal microbiota as described in Table S1. Cultivation of bacteria was carried out in laboratories of Ghent University and Vrije Universiteit Amsterdam – see below. The results from cultivation at the VU Amsterdam laboratory are depicted in Figure 1, 2, 3, 5, S1 and S2. The results from cultivation experiments at Ghent University are depicted in Figure 4 and S3. All strains were anaerobically cultivated from −80°C glycerol stocks (VU Amsterdam: NYCIII medium + 20% glycerol, Ghent: TSB + 15% glycerol) on agar plates under conditions indicated in Table S1. From these plates, colonies were picked to inoculate liquid New York City (NYC)-III broth with 10% horse serum as described previously (Hertzberger et al., 2022) or YEPG broth for *Candida albicans*. All cultures were handled under anaerobic conditions (VU Amsterdam: N_2_ + 10% CO_2_, Ghent University: 10% H_2_, 10% CO_2_, 80% N_2_) in an anaerobic chamber (VU Amsterdam: PLAS Labs 855 NB; Ghent university: Bugbox Plus).

**Figure 1:**
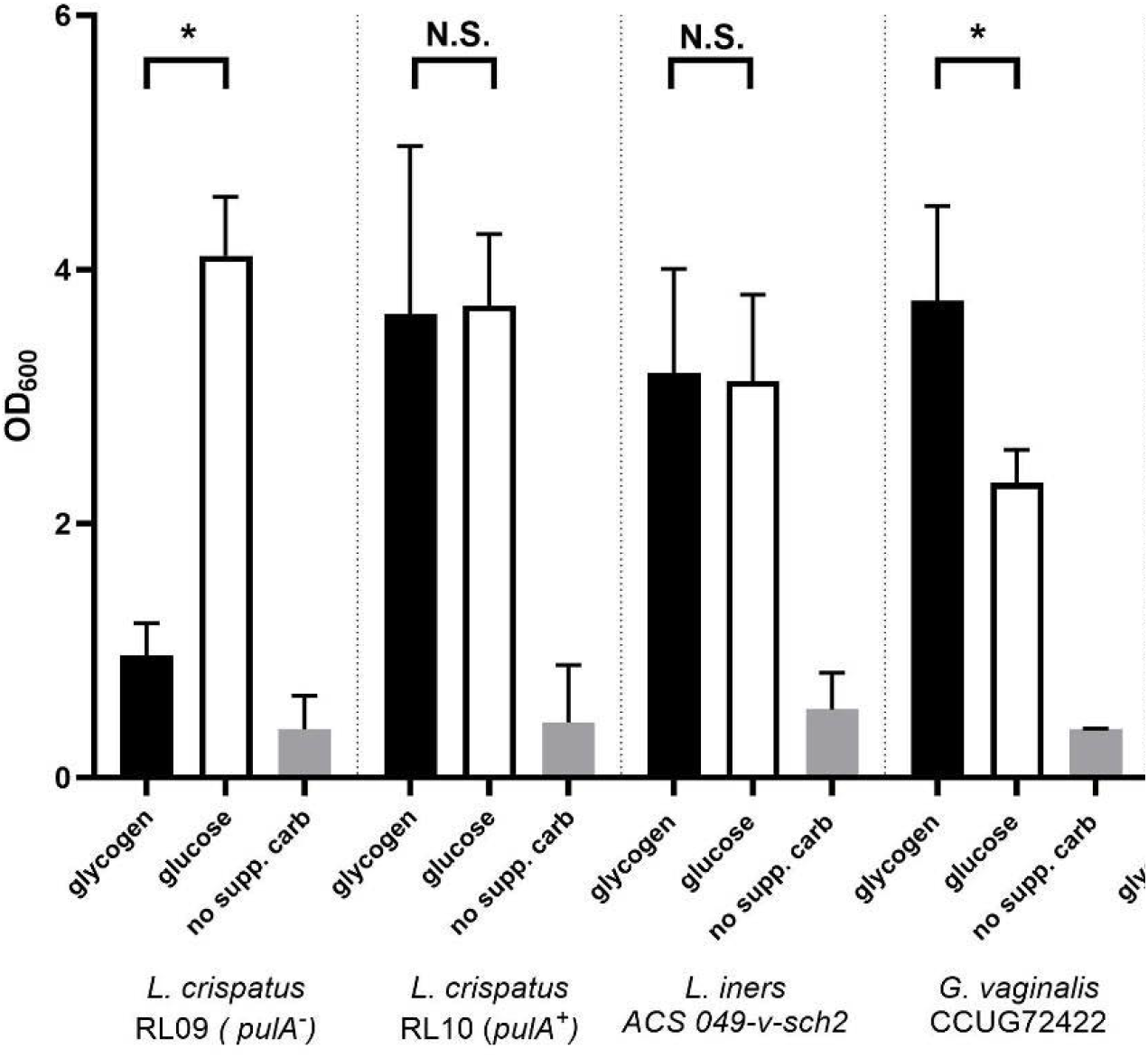
Optical density at 600 nm after 48 hours of growth in liquid NYCIII medium with 5 g/L glucose (white bars), 5 g/L glycogen (black bars) or no supplemented carbohydrate source (grey bars). All measurements were performed in three biological replicates in separate experiments on separate days. Data represent average and error bars are the standard deviation. Paired t-test was used to compare averages between glycogen and glucose. * = *p* < 0,05; N.S. = not significant

**Figure 2:**
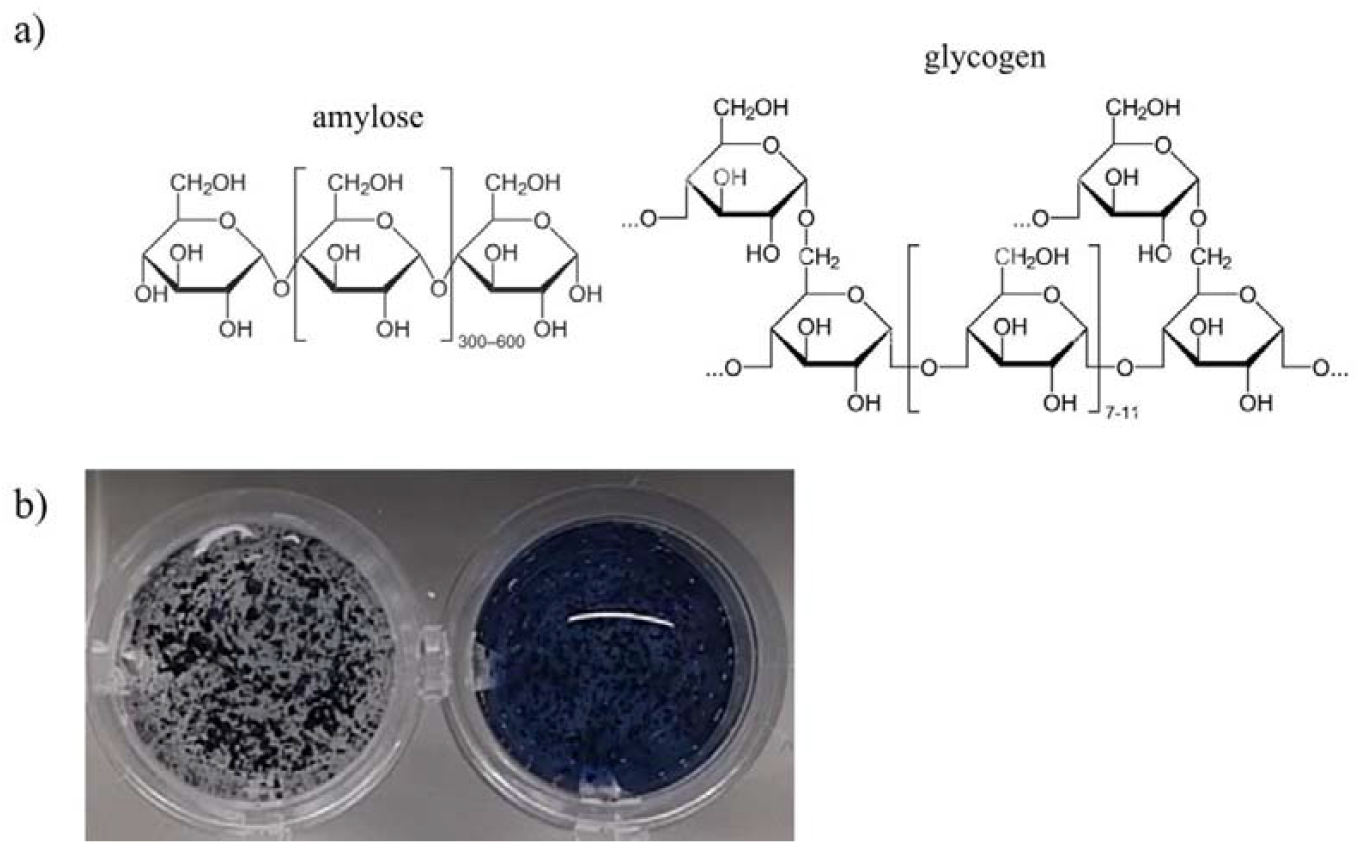
a) Comparison between amylose and glycogen structure; b) The AZCL-amylose substrate in xanthan gum buffer with buffer (negative control, left) or after mixing with saliva (positive control, right).

**Figure 3:**
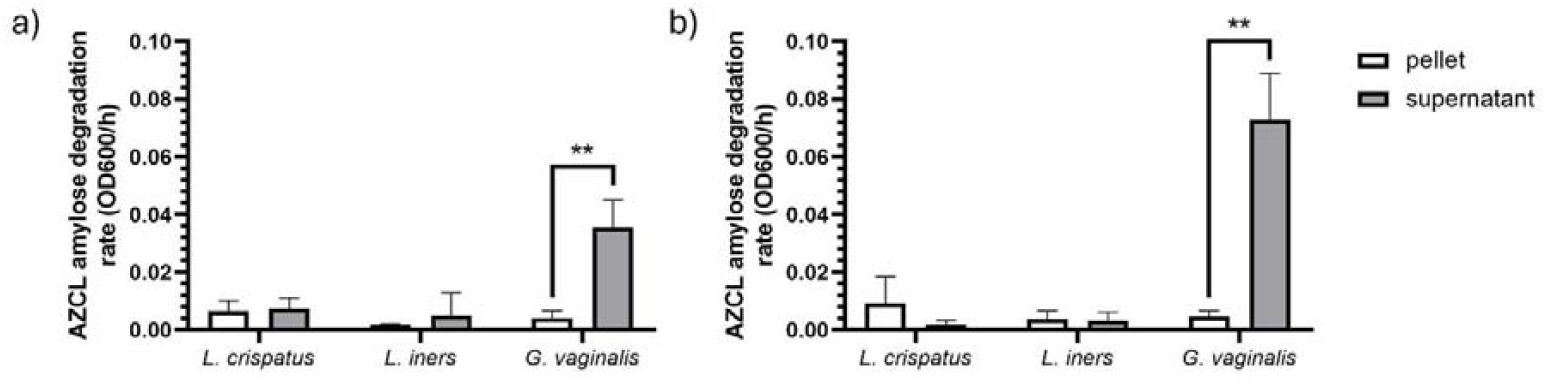
AZCL-amylose degradation rate expressed as the increase of optical density at 600 nm per hour in resuspended cell pellets (white bars) and supernatants (grey bars) corrected for optical density from *Lactobacillus crispatus* RL10 (*pulA*^+^), *Lactobacillus iners* ACS 049-v-Sch2 and *Gardnerella vaginalis* DSM 4944. Cultures were grown anaerobically in NYCIII medium with glycogen (panel a) or glucose (panel b) for 48 hours. Bars represent the mean and error bars represent standard deviation of three independent biological replicates. Pellets and supernatants were compared using t-test. ** = *p* < 0.005.

**Figure 4:**
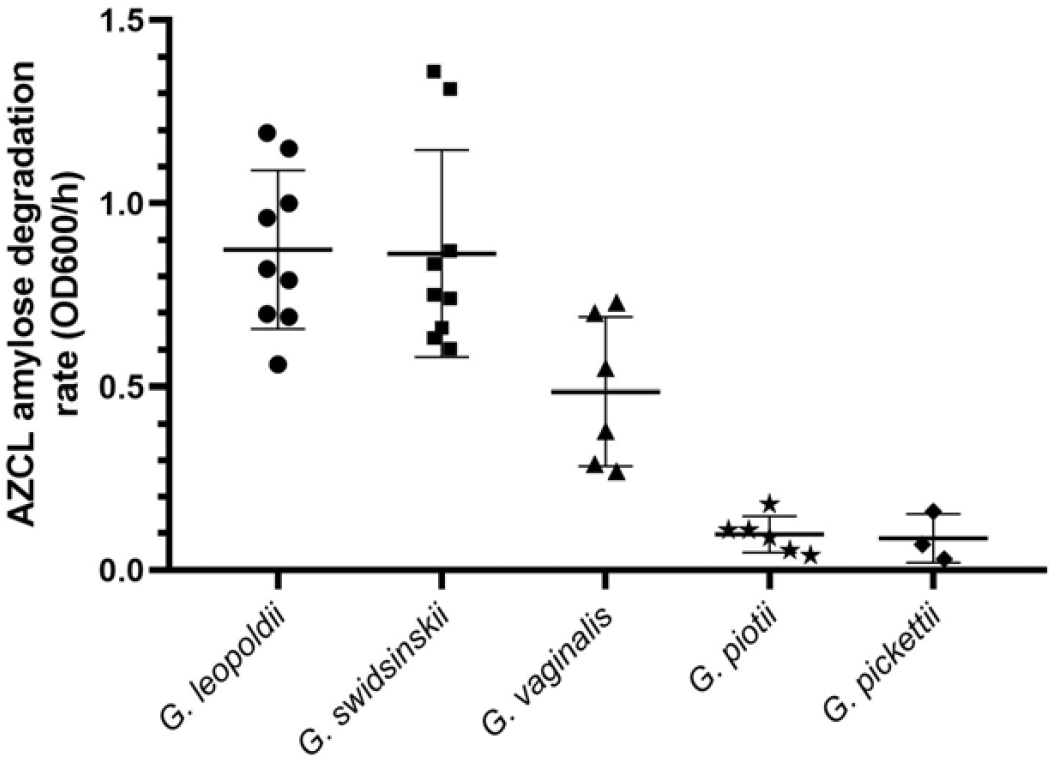
AZCL-amylose degradation rate by optical density increase at 600 nanometer of cultures (cells in growth medium) of *Gardnerella* species isolated by human vaginal samples grown for 48 hours on NYC III medium with glucose. The degradation rate was corrected for cell density (OD600). Data points represent biological replicates.

**Figure 5:**
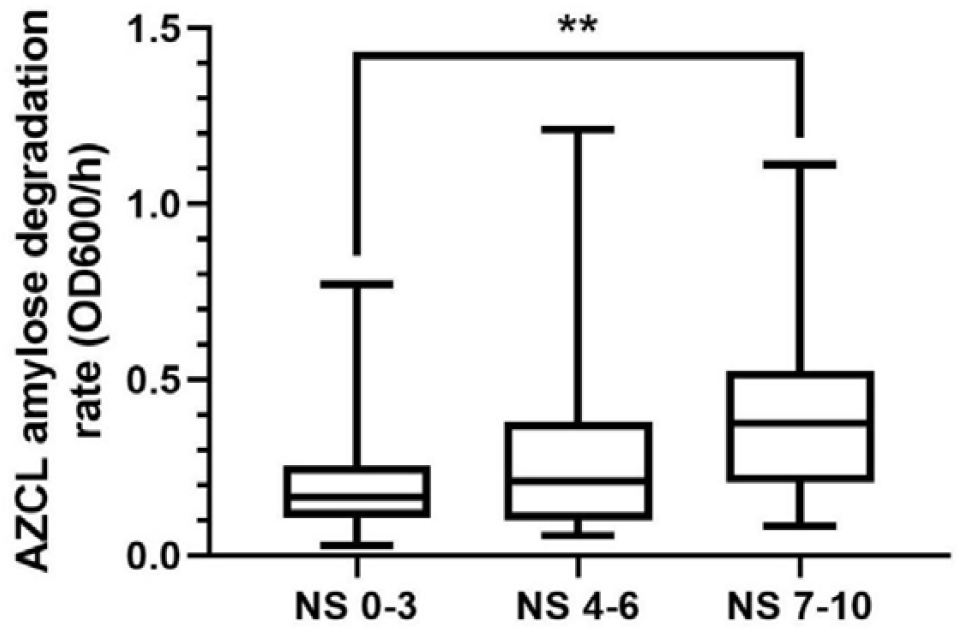
AZCL-amylose degradation rate of 96 vaginal samples with *Lactobacillus*-dominant microbiome (Nugent score 0-3), intermediate microbiome (Nugent-score 4-6) or bacterial vaginosis (Nugent score 7-10). *p*-values were calculated by ANOVA Tukey’s test.** = p < 0.005. Horizontal bars indicate the mean.

For three common vaginal bacterial species-*Gardnerella vaginalis, Lactobacillus iners* and *Lactobacillus crispatus*-we studied growth on glycogen by preparing a 1.1x New York City (NYC)-III broth with 10% horse serum to which we added a stock solution of 50 g/L oyster glycogen (Alva Aesar, Haver Hill, MA, USA) or 55 g/L glucose-monohydrate (Sigma-Aldrich) to obtain a final concentration of 5 g/L glucose-equivalents. We compared optical density after anaerobic incubation in NYCIII medium on these media. To verify whether any other medium components resulted in growth we inoculated these species also in NYCIII medium without glucose or glycogen supplemented. In Amsterdam, the optical density of cultures after 48 hours of growth was determined by diluting in saline and measuring the turbidity at 600 nm with a 1 mL cuvette using a Varian Cary 50 Bio UV-Vis Spectrophotometer (Spectra Lab Scientific Inc, ON, Canada), subtracting the cell-free medium control. In Ghent, the optical density was measured by diluting the culture in phosphate-buffered saline in a 96-well microtiter plate and measuring the turbidity at 600 nm in a GloMax® Explorer Multimode microplate reader (Promega, Madison, WI, USA).

### The AZCL-amylose assay

High purity dyed and crosslinked insoluble AZCL-Amylose (product code O-AZAMYF; Megazyme Inc., Bray, Ireland) was used as substrate for the assay of α-amylase. The AZCL-Amylose was used in a raw un-gelatinized form and suspended in an amylase buffer with xanthan gum to disperse the granules (Figure S1). After mixing with the sample (either vaginal sample of bacterial culture, spent medium or resuspended cell pellet) the increase in optical density in a microtiter plate was measured at 600 nm at 37°C. The lack of heating or other pretreatment results in the substrate retaining a coarse, granular, undissolved structure. Pelleting was prevented in this assay by suspension in a viscous buffer containing xanthan gum. Xanthan gum (Sigma-Aldrich Life Corporation, Saint Louis, MI, USA) was dissolved at 5 mg/mL in amylase buffer (100 mM sodium acetate buffer containing 5 mM CaCl_2_, pH adjusted with HCl to 5,3). This solution was autoclaved at 121°C for 20 minutes and kept at room temperature. AZCL-amylose is commercially available in two forms: regular and fine. The granules in the ‘fine’ product are small enough to pipet with regular pipet tips to dose and mix with the sample. This solution was stable without sedimentation of the raw amylose granules for up to 48 hours. AZCL-amylose was dispersed in the xanthan gum/amylase buffer prior to each experiment by shortly vortexing. The assay was carried out in a 96-well plate covered with adhesive film. A total of 190 µL reagent solution was added to 10 µL sample and mixed by carefully pipetting up and down. The microplate was incubated at 37 °C for 24 hours and absorption was measured at 600 nm every 10 minutes. Maximum rate was taken as the maximum slope of a three-hour time frame (including 19 data points). This rate was divided by the optical density at 600 nm of the culture. The enzyme assay was validated using *Aspergillus oryzae* amylase (Sigma-Aldrich) solution in amylase buffer with 150 units/µL where one unit corresponds to the amount of enzyme needed to cleave 1 µmol of maltose per minute at pH 6.0 and 25°C.

To study the prevalence of raw amylose degradation activity amongst *Gardnerella* species, we screened a set of 14 human vaginal *Gardnerella* strains from the species *G. leopoldii, G. swidsinskii, G. vaginalis, G. piotii* and *G. pickettii*. We compared these to strains of other common species colonizing the human vagina including *Candida albicans, Fannyhessea vaginae* (previously known as *Atopobium vaginae*), *Prevotella bivia, Bifidobacterium* and several *Lactobacillus* species. Cultures were either used without centrifugation or centrifuged for 1 minute at 20,000 rcf at room temperature to separate cells. Spent medium was aspirated and mixed with the substrate and cell pellets were resuspended in sterile phosphate-buffered saline and mixed with the substrate. The degradation rate was divided by the optical density at 600 nm.

### The AZCL-amylose assay and quantification of bacteria in vaginal swabs

Vaginal swabs were collected from patients visiting the Center for Sexual Health (Amsterdam, The Netherlands), as described previously (Dols et al., 2016). The swabs were eluted in Amies buffer and 15% glycerol and cysteine was added for cryoprotection to preserve live bacteria and enzymes for subsequent isolation and analysis. The samples were stored at −80°C. Samples for this study were selected on the basis of Nugent scores choosing one third of samples with scores between 0-3 (healthy), one third with scores between 4-6 (intermediate) and one third with a Nugent score between 7-10 (bacterial vaginosis). For the amylose assay, samples were thawed, vortexed and pipetted but not homogenized or centrifuged, which means that mucus and other solid material was still present in the sample when pipetted. Of the liquid fraction, 10 µL was transferred to a microtiter plate to which 190 µL of the AZCL-amylose in the amylase buffer with xanthan gum was added. The assay and rate calculation was carried out as described above.

The commercially available multiplex PCR for ATRiDA test BV diagnosis (ATRiDa, Amersfoort, The Netherlands) was performed according to instructions of the manufacturer ((Rumyantseva et al., 2016; Shipitsyna et al., 2020) on isolated DNA from vaginal samples targeting *Lactobacillus* spp., total *Gardnerella*, total *Fannyhessea* and total bacteria. Both the DNA extraction as well as the qPCR reaction were performed as described previously (van der Veer et al., 2018). The same DNA extracts were utilized for species specific *G. vaginalis, G. piotii, G. leopoldii* and *G. swidsinskii* qPCR assays (Latka et al., 2022). To account for multiple testing, we adjusted the *p*-value threshold by Bonferroni-correction leading to a required *p*-value of 0.013 (*p*=0.05 divided by four outcomes) for the BV diagnostic PCR and the *Gardnerella* species-specific qPCR. We used the log transformed data to find correlations between AZCL-amylose degradation rate and abundance of any of the targets. All data processing and statistical analysis were performed in Microsoft Excel and GraphPad Prism 9.5.0.

## Results

### *Gardnerella* and *Lactobacillus iners* can use glycogen as a carbon source for growth

The optical density of *L. crispatus, L. iners* and *G. vaginalis* after anaerobic growth in NYCIII medium with glucose or glycogen is shown in Figure 1. As expected, and previously shown (Hertzberger et al., 2022; van der Veer et al., 2019), the amylopullulanase-deficient strain of *L. crispatus* did not grow on glycogen, whereas the amylopullulanase-sufficient strain grew to a comparable optical density on glycogen as on glucose. The *L. iners* strain showed an equal optical density on glycogen compared to glucose, while the *G. vaginalis* strain showed a higher optical density on glycogen compared to glucose. To verify that the increase in turbidity was solely due to growth on the carbohydrate source added (glucose or glycogen) we added a control–the NYC medium without any glucose or glycogen. No increase in optical density was observed in this medium compared to the cell-free control (Figure 1).

### Multi-well assay with undissolved labeled amylose to measure enzymatic rate of α-glucosidases

Next, we further characterized α-glucosidase activity of *L. crispatus, L. iners* and G. *vaginalis* using a quantitative amylase assay with raw amylose labelled with a covalently linked azure cross-linked (AZCL) chromogenic substrate (McCleary, 1980) in a microtiter plate. This chromogenic substrate is conventionally used to study amylase activity using agarose plates or in dissolved form but is applied here using undissolved granules in a xanthan gum buffer to keep the substrate dispersed. The release rate of blue dye is dependent on the enzymatic digestion rate of the labelled glycan (see calibration curve in Figure S2). No pretreatment, purification or dialysis of the sample is required, and the assay is suitable for use in multi-well format in a plate reader as well as for visual detection.

### *Gardnerella vaginalis* shows extracellular amylose-degrading activity

In Figure 3, the enzymatic activity in pellets and supernatants as determined by the amylose degradation assay is shown. The amylose degradation assay showed that the investigated *G. vaginalis* strain was capable of degrading the raw amylose substrate, while in *L. crispatus* and *L. iners* cultures, no labeled amylose digestion was observed. The activity was mostly found in supernatants of *G. vaginalis* compared to the cell pellet (*p*=0.005). The activity was also present in *G. vaginalis* culture grown on medium with glucose, indicating that *G. vaginalis* α-glucosidase activity is not subject to carbon-catabolite repression, as observed for the glycogen-degrading activity of *L. crispatus*. Similarly to what was observed in glycogen growth, also for glucose the supernatant was more active compared to the pellet (*p*= 0.002).

### *Gardnerella* species show variation in AZCL-amylose degradation rate

AZCL-amylose degradation activity of different *Gardnerella* species is shown in Figure 4. Overall, the data obtained indicate a clear difference in the ability to degrade amylose among certain *Gardnerella* species. Using Tukey’s test for multiple comparisons, *G. leopoldii* showed highly significant differences compared to *G. piotii* (*p* < 0.0001) and *G. pickettii* 3 (*p* < 0.0001), with mean differences in amylose degradation rates of 0.776 and 0.787, respectively. Similarly, *G. swidsinskii* exhibited a significant difference compared to *G. piotii* (*p* < 0.0001) and *G. pickettii* (*p* < 0.0001). On the other hand, some comparisons were not statistically significant, such as between *G. leopoldii* and *G. swidsinskii* (*p* = 0.99) and between *G. vaginalis* and *G. pickettii* (*p* = 0.08).

### AZCL-amylose degradation rate is associated with Nugent score

The substantial differences in amylose degradation activity between three of the most common vaginal species prompted us to study the presence of this enzymatic activity in vaginal swabs. The degradation rate in clinical samples across Nugent categorization is shown in Figure 5. Degradation of amylose was strongly associated with Nugent scoring. Samples from women with Nugent score between 7-10 (bacterial vaginosis) showed a mean AZCL-amylose degradation rate of 0.41 OD_600_/h +/-0.27 compared to 0.22 OD_600_/h +/-0.18 in women with Nugent score 0-3 (*p*=0.0051). The results of the qPCR for the genera *Gardnerella, Fannyhessea* and total bacteria were strongly associated with Nugent score. These species had a higher abundance in samples with Nugent score 7-10 compared to samples with a Nugent score of 0-3, while lactobacilli were more abundant in women with low Nugent scores compared to women with high Nugent scores. All correlations between the genera and Nugent score group are shown in Table S2.

In growth experiments, a higher AZCL-amylose degradation rate was observed in cultures of *Gardnerella* species compared to *Lactobacillus* species. Therefore, we hypothesized that the observed difference in samples from women with high Nugent scores compared to women with low Nugent scores may be due to higher *Gardnerella* abundance in the high Nugent score group. We therefore assessed whether the measured amylose degradation rate was associated with *Gardnerella* abundance, and found a positive correlation between AZCL-amylose degradation rate and levels of *Gardnerella* and *Fannyhessea* (Spearman correlation of 0.24 and 0.23, *p*-value of 0.016 and 0.028, respectively). The highest degradation rates were found in samples containing multiple *Gardnerella* species, including *G. leopoldii*, G. *swidsinskii*, and *G. piotii* (Figure S3).

### AZCL-amylose degradation and sialidase activity in *Gardnerella* species

Four strains of *Gardnerella*, each representing a distinct species, were selected for comparative analysis. After 48 hours of growth, culture supernatants were harvested to assess AZCL-amylose degradation and sialidase activity. *G. swidsinskii* and *G. leopoldii* demonstrated higher AZCL-amylose degradation activity relative to *G. vaginalis* and *G. piotii*. Notably, pronounced sialidase activity was observed only in *G. piotii* strain 1801, consistent with the presence of the *nanH3* gene (Kurukulasuriya et al., 2021)(14). *G. vaginalis* exhibited intermediate sialidase activity (Figure 6).

**Figure 6:**
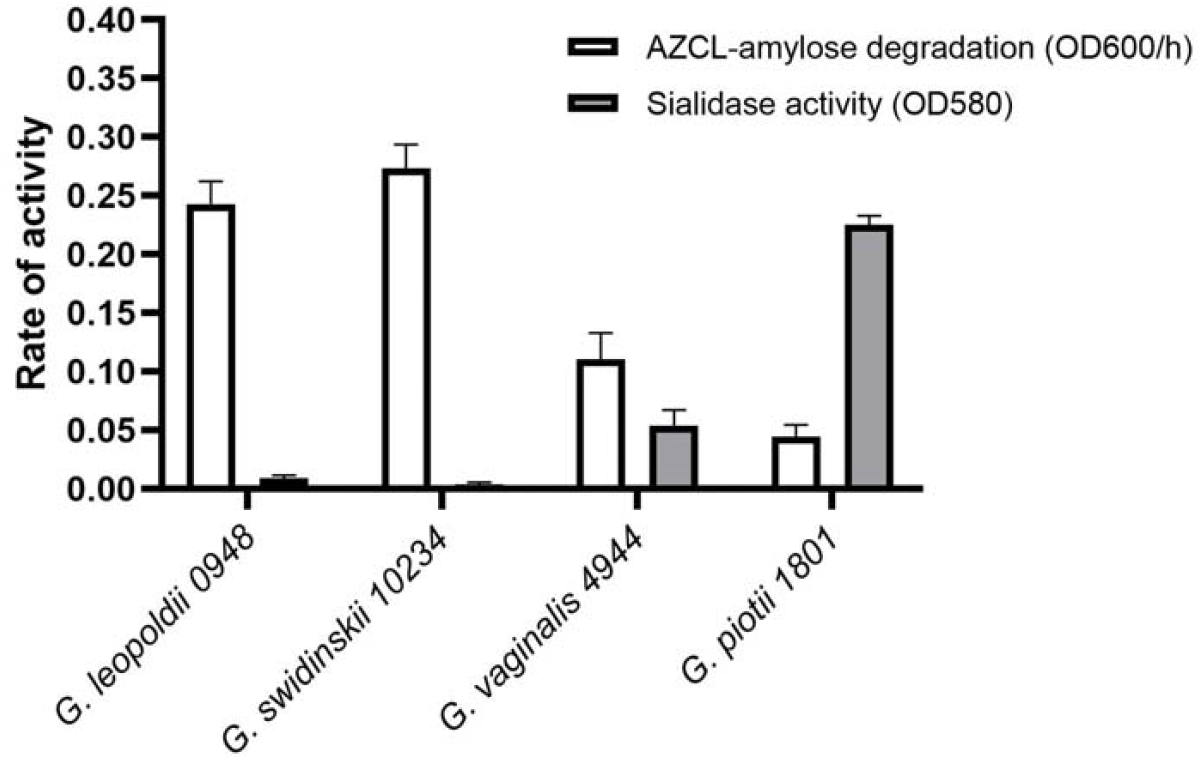
Amylose degradation rates and sialidase activity in supernatants of four *Gardnerella* species.

## Discussion

In this study, we investigated the presence of extracellular α-glucosidase activity among key members of the vaginal microbiota. These enzymes catalyze the breakdown of α1–4 and/or α1–6 glycosidic linkages in α-glucans such as glycogen, amylopectin, and amylose. Between puberty and menopause, the female reproductive tract contains substantial glycogen reserves (Mirmonsef et al., 2014), which may serve as a carbon source for bacteria that express and secrete glycogen-degrading enzymes. Previously, we demonstrated that *Lactobacillus crispatus* strains can degrade starch and glycogen (Hertzberger et al., 2022; van der Veer et al., 2019). Jenkins et al. identified and characterized the amylopullulanase responsible for this activity in *L. crispatus* (Jenkins et al., 2023). However, in the current study, we found that *L. crispatus* was unable to degrade raw amylose—composed primarily of α1–4 linkages—whereas *Gardnerella* species demonstrated a clear capacity to do so. Previously, resistant starches where found to modulate the gut microbiome (I. Brown et al., 1997; Jung et al., 2020; Maier et al., 2017; Sybille et al., 2013). High amylose corn starch is degraded by specific gut genera, one of which is *Bifidobacterium* that is closely related to the genus *Gardnerella*. The amylopullulanase of bifidobacteria has the same carbohydrate-binding, amylase and pullulanase domains, N-terminal signal peptide and C-terminal transmembrane domain as the *Gardnerella* enzyme that is likely responsible for the observed raw amylose degradation activity in this study (Bhandari et al., 2023; Jenkins et al., 2023).

Interestingly, we observed clear differences between *L. crispatus* and *G. vaginalis* α-glucosidase activity not only in substrate specificity but also in its regulation and localization. In contrast to *L. crispatus*, which exhibits catabolite repression of amylopullulanase in the presence of glucose and maltodextrins (Hertzberger et al., 2022), *Gardnerella vaginalis* retained α-glucosidase activity even when grown on glucose. Additionally, while *L. crispatus* amylase activity is largely cell-associated, the *Gardnerella* activity was primarily extracellular, despite the presence of a C-terminal transmembrane domain. It is possible that the physical structure of our chemically labelled amylose substrate limited access for cell-bound enzymes, or that proteolytic cleavage of the transmembrane domain releases the enzyme into the environment. This phenomenon mirrors what has been described for *Gardnerella* sialidase, which is also partially released into culture supernatants despite having a membrane anchor (Robinson et al., 2019). Future experiments are required to precisely localize these enzymes and investigate post-translational cleavage.

Although AZCL-amylose degradation activity correlated well with Nugent scores, we did not observe a clear association with the abundance of any single *Gardnerella* species. For example, *G. swidsinskii*–dominant samples exhibited both high and low degradation activity, and several high-activity samples lacked detectable *G. swidsinskii* or *G. leopoldii*. However, the highest degradation rates were found in samples containing multiple *Gardnerella* species (*G. leopoldii, G. swidsinskii*, and *G. piotii*). These findings suggest strain-level variation within species, and potential cooperative interactions when multiple species are present. While host-derived amylase has been detected in vaginal fluid and proposed to contribute to glycogen degradation (Nasioudis et al., 2015; Nunn et al., 2020; Spear et al., 2014), our data suggest that its role may be limited in *Lactobacillus*-dominated communities, given the generally low amylose degradation activity in samples with Nugent scores of 0–3. We cannot exclude a contribution of human amylase in BV-associated communities, but the absence of detectable degradation by other BV-associated genera (e.g., *Fannyhessea, Prevotella*, and vaginal *Bifidobacterium*) leaves *Gardnerella* as the most likely microbial source.

To contextualize *Gardnerella*’s α-glucosidase activity within a broader enzymatic profile, we also measured sialidase activity. Our data suggest division of labor among species: *G. swidsinskii* and *G. leopoldii* exhibited higher amylose-degrading activity compared to *G. vaginalis* and *G. piotii*, while *G. piotii* was the only strain with notable sialidase production. This is consistent with prior reports linking bacterial vaginosis (BV) not to specific *Gardnerella* species but rather to increased species diversity within the genus (Munch et al., 2024). While our data suggest a potential trend between amylase and sialidase activity across species, the limited sample size precludes any definitive conclusions regarding an inverse correlation.

Taken together, the characteristics of the *Gardnerella* α-glucosidase could explain the low glycogen levels in samples with BV (Mirmonsef et al., 2014). Lactobacilli may in contrast only ‘graze’ the accessible linkages of the glycogen molecules and cease amylopullulanase expression as soon as the smaller maltodextrins start to accumulate (Hertzberger et al., 2022). Previously, *Gardnerella* sialidase and cytolysin were found to be linked to clinical characteristics of BV, namely mucus thinning and epithelial exfoliation (Gilbert et al., 2013; Lewis et al., 2013). Here, we propose that *Gardnerella*’s glycogen metabolism represents a third clinical characteristic of BV. This activity links its colonization and overgrowth to the reduced glycogen levels observed in the vaginal lumen during BV. Understanding the enzymatic capabilities and interspecies variation of *Gardnerella* may be crucial to further unravel its role in BV pathophysiology.

The ability of *Gardnerella* species to secrete extracellular enzymes that degrade resistant glycogen structures such as amylose highlights a potential microbial mechanism contributing to glycogen depletion in BV. These findings suggest that targeting bacterial α-glucosidase activity—specifically the amylase-like enzymes of *Gardnerella*—could represent a novel therapeutic strategy for the prevention or treatment of BV. Inhibiting this activity may help preserve vaginal glycogen levels, thereby supporting the growth of beneficial lactobacilli and promoting a more stable, protective microbial community. Future work should explore the feasibility and specificity of such targeted interventions.

## Acknowledgments

VU Amsterdam: Alexander Woudstra, Ritesh Panchoe for experiments setting up anaerobic cultivation. Jurgen Haanstra for supervision of students. Douwe Molenaar for help with data analysis and statistics. Martin Braster for help with the anaerobic chamber.

GGD Amsterdam: Mirjam Dierdorp for isolating the DNA from vaginal samples.

Ghent University: Mario Vaneechoutte for providing different vaginal strains. Leen van Simaey and Aliona Rosca for their assistance in AZCL-amylose assays of different species of the vaginal microbiome.

Megazyme kindly donated 5 grams of fine AZCL-amylose for this study.

## Author contributions

Conceptualization, R.H., R.K. and W.L.; Methodology and data analysis, R.H., L. H., L. van S., W.L., R.K., P.C., S.B., S.M.; Experiments: R.H., L. H., L. van S., S.M.; Data curation: R.H., L.H., L. van S.; Writing—original draft preparation, R.H.; Writing review and editing, R.H., L.H., S.B., P.C., S.M., R.K.; Supervision: S.B., P.C., R.K. All authors have read and agreed to the published version of the manuscript.

