## Supplemental File for "Degradation of resistant α-1,4-glucan by vaginal *Gardnerella* species is associated with bacterial vaginosis"

**Supplementary material**

This supplementary material is associated with the paper by Hertzberger et al. (2025). Degradation of resistant α-1,4-glucan by vaginal *Gardnerella* species is associated with bacterial vaginosis.

**Table S1** Strains used in this study and the agar plates that were used for preculturing. CA, chocolate agar; CNA, Columbia Naladixic Acid agar; TSA+5%SB, tryptic soy agar + 5% sheep blood; NYCIII, New York City agar + 10% horse serum.

| Species | Strains | Agar plate medium and incubation conditions |
| --- | --- | --- |
| *Bifidobacterium breve* | FB034-05AEa | 37°C, anaerobic, TSA+5%SB |
| *Candida albicans* | ATCC 90028 | 30°C, aerobic, Sabouraud |
| *Fannyhessea vaginae* | ACS-043-V-Col2 | 37 °C, anaerobic, CNA |
| *Gardnerella pickettii* | CCUG 39787 | 37 °C, anaerobic, CA |
| *Gardnerella leopoldii* | UGent 06.41^T^ = CCUG 72425 | 37 °C, anaerobic, CA |
|  | BV13.2.3.2.1 | 37 °C, anaerobic, CA |
|  | CCUG 28832 | 37 °C, anaerobic, CA |
|  | UGent 09.48 = CCUG 72426 | 37 °C, anaerobic, CA |
| *Gardnerella piotii* | UGent 21.28 | 37 °C, anaerobic, CA |
|  | UGent 18.01^T^ = CCUG 72427 | 37 °C, anaerobic, CA |
|  | CCUG 39708 | 37 °C, anaerobic, CA |
| *Gardnerella swidsinskii* | CCUG 7733 | 37 °C, anaerobic, CA |
|  | CCUG 31348 | 37 °C, anaerobic, CA |
|  | UM034 BV | 37 °C, anaerobic, CA |
| *Gardnerella vaginalis* | UGent 09.01 | 37 °C, anaerobic, CA |
|  | UGent 25.49 = CCUG 72423 | 37 °C, anaerobic,  Ghent: CA; Amsterdam: NYCIII agar |
|  | UGent 09.07 = CCUG 72422 | 37 °C, anaerobic,  Ghent: CA; Amsterdam: NYCIII agar |
| *Lactobacillus crispatus* | LMG 0479^T^ | 37 °C, anaerobic, NYCIII agar |
|  | RL10 | 37 °C, anaerobic, NYCIII agar |
|  | RL09 | 37 °C, anaerobic, NYCIII agar |
| *Lactobacillus gasseri* | BVS30A2 | 37 °C, anaerobic, CNA |
|  | FWOBV0322 | 37 °C, anaerobic, NYCIII agar |
| *Lactobacillus iners* | FB123-CNA-4 | 37 °C, anaerobic, NYCIII agar |
| *Lactobacillus jensenii* | FWOBV0433 | 37 °C, anaerobic, NYCIII agar |
| *Prevotella bivia* | ACS-045-V-Col2 | 37 °C, anaerobic, CA |

**Table S2**. Association between Nugent Score and average concentration gene equivalents (ACGE) by bacterial vaginosis molecular diagnostic qPCR (ATRiDa).

|  | ACGE at Nugent scores | | |  |
| --- | --- | --- | --- | --- |
| Genus | 0-3 | 4-6 | 7-10 | *p*-value (Mann-Whitney) Nugent groups 0-3 and 7-10 |
| *Gardnerella* | 6.6*10^5^ | 3.0*10^6^ | 1.3*10^7^ | <0.0001 |
| *Fannyhessea* | 1.7*10^6^ | 1.2*10^6^ | 3.3*10^7^ | <0.0001 |
| *Lactobacillus* | 7.5*10^7^ | 4.5*10^7^ | 3.7*10^7^ | 0.0026 |
| Total bacteria | 7.6*10^7^ | 5.4*10^7^ | 2.4*10^8^ | <0.0001 |

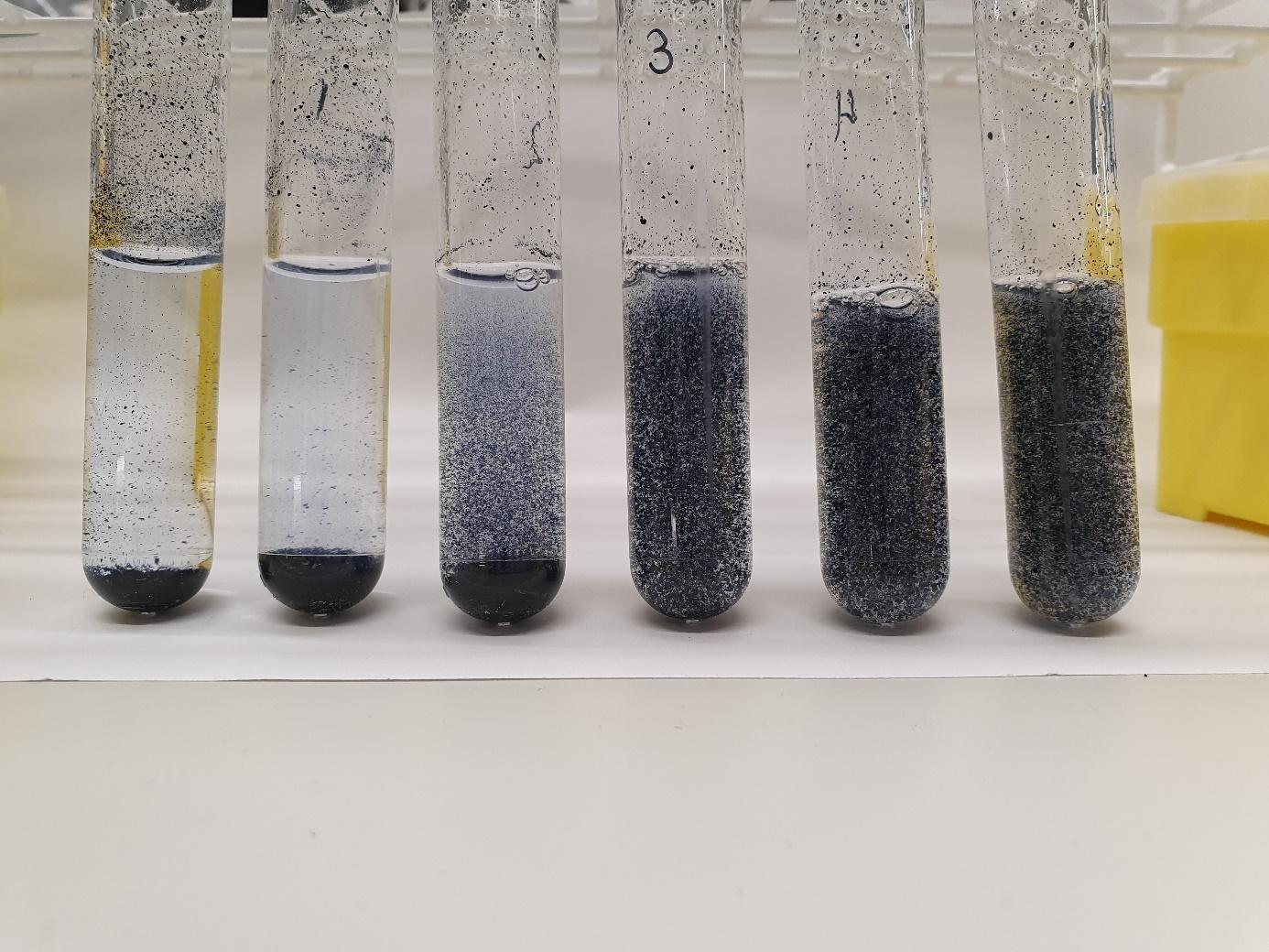

**Figure S1** Pelleting and dispersion of 2 mg/mL AZCL-amylose granules at different xanthan gum concentrations (0-5 mg/mL) in amylase buffer (pH 5,3) after 48 hours.

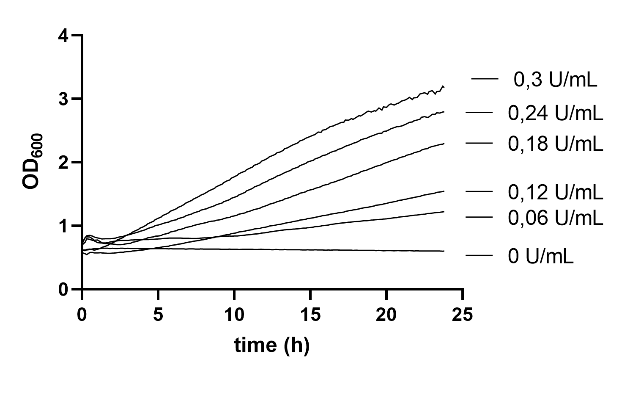

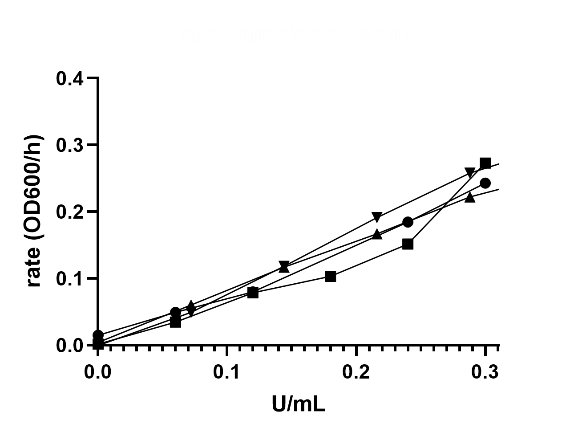

**Figure S2**. AZCL-amylose degradation rate by optical density increase at 600 nm of full cultures (cells in growth medium) of common bacterial and fungal members of the vaginal microbiome grown for 48 hours on NYC III glucose or YEPG (*Candida albicans*). Data points represent biological replicates from independent experiments that were corrected for cell density. The Y-axis has been adjusted to allow comparison with amylose degradation of the *Gardnerella* strains in Figure 4.

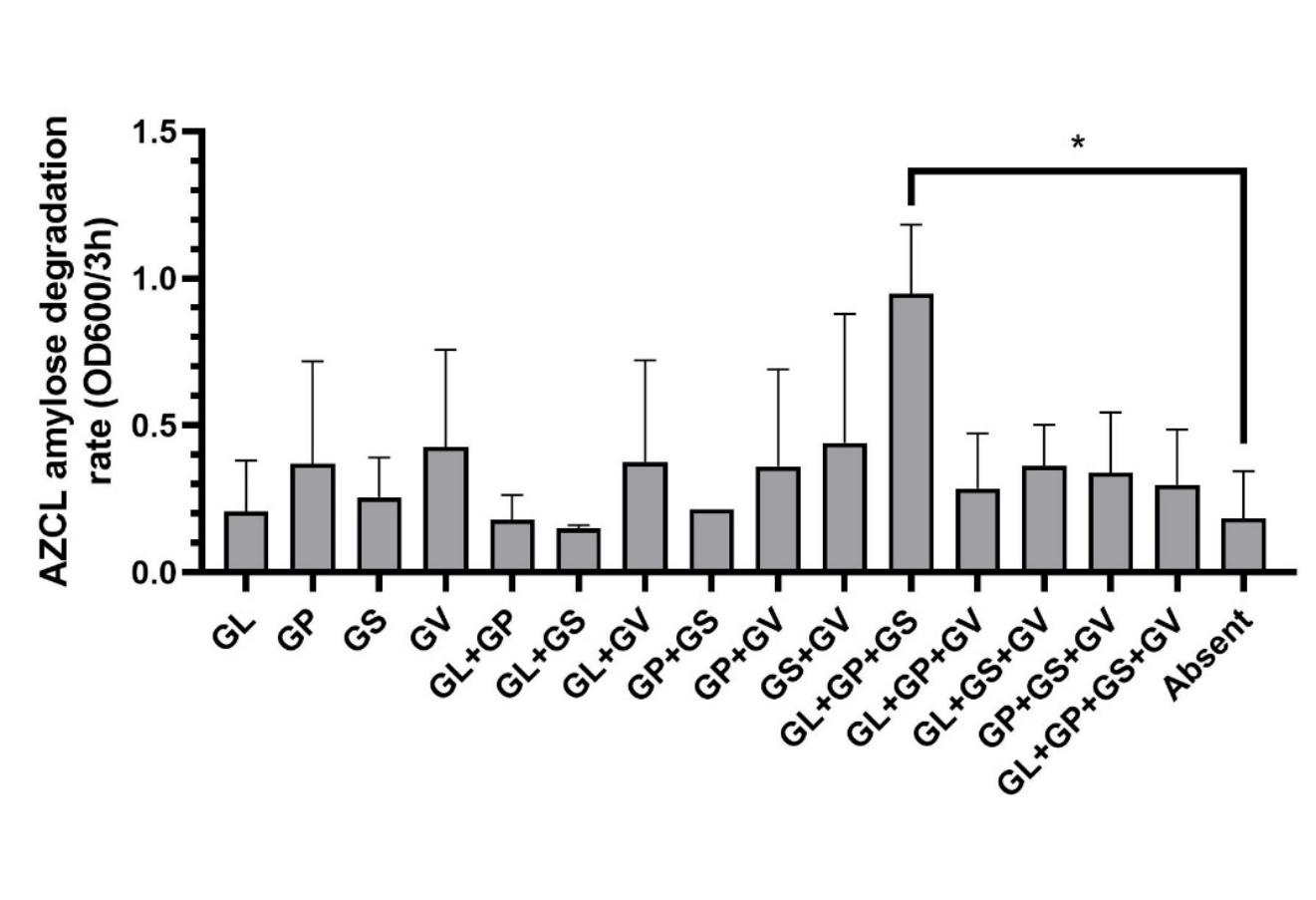
**Figure S3.** AZCL-amylose degradation rate measured by the increase in optical density at 600 nm in vaginal samples classified by the presence of *Gardnerella* species as detected by PCR. *GL: G. leopoldii; GS: G. swidinskii; GV: G. vaginalis; GP: G. piotii. p*-values were calculated by ANOVA Tukey's test.* *p* < 0.05.
